# Developmental changes in the control of primary motoneuron excitability by the M-current in larval zebrafish

**DOI:** 10.1101/2024.10.17.618843

**Authors:** Stephanie F. Gaudreau, Tuan V. Bui

## Abstract

Spinal circuits for locomotion undergo maturation during early development. How intrinsic properties of individual spinal neuron populations change throughout motor maturation is not fully understood. Here we identify for the first time the presence of the persistent outward potassium current known as the M-current in primary motoneurons of larval zebrafish. We show that the M-current controls excitability of primary motoneurons and its role in excitability control changes during development such that the magnitude of the M-current in primary motoneurons transiently increases at 3 days post-fertilization. These findings reveal a novel mechanism by which control over excitability of primary motoneurons in larval zebrafish is ensured, underscoring developmental changes in ion current contributions to intrinsic properties. Broadly, these data support the M-current as a conserved means to control motoneuron excitability across vertebrates.

## INTRODUCTION

In early development, vertebrates transition through a series of progressively more complex movements. Eventually, underlying spinal locomotor circuits become sufficiently refined to produce locomotion. The precise changes that spinal locomotor circuits undergo during development are still not fully understood. Amongst several possibilities, spinal neurons can undergo changes in synaptic receptors (Laliberte et al., 2022; Wilson et al., 2004) and ion currents (Sharples and Miles, 2021) during development.

The developing zebrafish (*Danio rerio*) is an attractive vertebrate model for studying how the spinal cord mediates locomotion. Larval zebrafish have a relatively simple nervous system such that spinal neurons are easily identifiable and accessible for study (Bernhardt et al., 1990; Myers et al., 1986). The larval zebrafish is especially well-suited to the study of motor control maturation because of their rapid development and the stereotyped emergence of motor behaviours during this time (Budick and O’Malley, 2000; Drapeau et al., 2002). As of 17 hours post-fertilization (hpf), zebrafish embryos produce large amplitude body bends known as coiling (Kimmel et al., 1995; Saint-Amant and Drapeau, 1998). Soon after, around 24 hpf, they can execute consecutive coils to either side of the head, referred to as double coiling (Knogler et al., 2014). At 3 days post-fertilization (dpf), zebrafish larvae exhibit their first instances of spontaneous swimming (Saint-Amant, 2010). The swimming is erratic and long-lasting but infrequent. This swimming rapidly evolves into what is known as the beat-and-glide swim by 4 dpf (Budick and O’Malley, 2000; Buss and Drapeau, 2001). This mature form of larval zebrafish locomotion consists of more controlled, low-amplitude rhythmic tailbeats interspersed by periods of gliding. These stereotyped transitions in locomotion make it possible to link changes in motor output to changes in spinal circuits, ultimately shedding light onto the mechanisms underlying the spinal control of movement (Roussel et al., 2021, 2020).

Major changes to spinal locomotor circuits of developing zebrafish have been identified including the addition of new neurons (Myers, Eisen, and Westerfield 1986; Kimmel et al. 1995; Kuwuda et al. 1990; Satou, Kimura, and Higashijima 2012), progressive addition of chemical neurotransmission to a gap-junction-prominent neural scaffold (Knogler et al., 2014; Roussel et al., 2020; Saint-Amant and Drapeau, 2001), formation and refinement of synaptic connections (Bernhardt et al., 1990; Liu and Westerfield, 1988; Roussel et al., 2021) as well as incorporation of neuromodulatory systems (Brustein et al., 2003; Jay et al., 2015; Lambert et al., 2012; Montgomery et al., 2018; Thirumalai and Cline, 2008). Even the firing of spinal neurons changes through development (Buss et al., 2003; Saint-Amant and Drapeau, 2000) though the ion currents underlying these changes remain to be fully identified.

Primary motoneurons (pMNs) are the largest and earliest born motoneurons in larval zebrafish (Myers, 1985). They control the fastest and largest amplitude movements (Liu and Westerfield, 1988). These types of movements are most prominent in zebrafish 2 dpf or younger and become less prominent as of the third day of development. By this age, swimming involves secondary motoneurons (sMNs) with the involvement of pMNs becoming increasingly less prominent (Liu and Westerfield, 1988; McLean et al., 2007). Recently, it has been demonstrated that different types of mouse motoneurons express varying magnitudes of the persistent subthreshold potassium current known as the M-current (Sharples et al., 2023). Sharples et al. demonstrate that the M-current expression in motoneurons helps control their recruitment (Sharples et al., 2023). Given that the M-current has been shown to work in tandem with the persistent sodium current in mice (Verneuil et al., 2020), and that zebrafish spinal neurons possess a persistent sodium current (Benedetti et al., 2016; Song et al., 2020), we posited that motoneurons in larval zebrafish may also express the M-current. Furthermore, we hypothesize that the role of the M-current in shaping pMN excitability may change throughout development as the most prominent locomotor movements rely increasingly less on pMNs with the emergence of secondary MNs that are relatively more involved in swimming (Ampatzis et al., 2014; McLean et al., 2007).

Using whole-cell patch-clamp electrophysiology, we reveal for the first time the presence of the M-current in larval zebrafish spinal circuits for locomotion. We demonstrate that the M-current regulates excitability of pMNs and show that the magnitude of the M-current in pMNs changes throughout development. Together, these data describe a novel mechanism of motoneuron excitability control in the larval zebrafish.

## METHODS

### Animal Care

All experiments were performed in accordance with the protocol approved by the University of Ottawa’s Animal Care Committee (BL-4416). Adult zebrafish are maintained at 28.5°C with a 14 hour on/10 hour off light cycle, with lights off at 11PM and on at 9AM. Embryos are fertilized between 9AM-10AM and stored in embryo medium at 28.5°C until use for experimentation at 2 to 5 dpf.

### Preparation for electrophysiology

Larval zebrafish are anesthetized in 0.02% tricaine (MS-222, Aqualife TMS; Syndel Laboratories) before being pinned down through the notochord onto a Sylgard (Dow Corning) coated dish. Pins are made using Tungsten wire (0.25 mm); one is placed caudally near the tip of the tail, and the other is placed rostrally near the center of the yolk sac. Spinalization is performed using fine surgical scissors at the level of segments 2-3. The skin is then peeled back between the two pins using fine forceps. Next, larvae are bathed in 1 mg/mL collagenase (Millipore Sigma; incubation times: 8 minutes at 2 dpf, 10 minutes at 3 dpf, 12 minutes at 4 dpf, and 15 minutes at 5 dpf) to facilitate muscle removal. The collagenase solution is rinsed out 4-5 times with artificial cerebrospinal fluid (aCSF) containing: 134 mM NaCl, 2.9 mM KCl, 1.2 mM MgCl_2_, 2.1 mM CaCl_2_, 10 mM dextrose, and 10 mM HEPES (pH of 7.8 adjusted with NaOH and osmolarity between 280-290 adjusted with sucrose). Muscle removal is performed using suction through a wide-bored glass capillary (typically, the tips are gently broken to get the right sized tip for muscle removal). Muscles overlaying the spinal cord are removed to expose the spinal cord for whole-cell patch-clamp electrophysiology.

### Whole-cell patch-clamp electrophysiology

Electrodes for whole-cell patch-clamp experiments are made using borosilicate glass capillaries (Sutter Instrument; BF150-110-10). Tip resistances ranging from 5 to 7 megaOhms (MOhms) were used. Intracellular recording solution used contained the following: 16 mM KCl, 116 mM K-gluconate, 4 mM MgCl_2_, 10 mM HEPES, 10 mM EGTA, and 4 mM Na2-ATP, adjusted to a pH of 7.2-7.3 with KOH. Slight positive pressure was applied during the descent toward the spinal cord to prevent debris from contaminating the tip of the electrode. This positive pressure was used to break the dura thereby exposing the neurons within the spinal cord. Positive pressure was decreased slightly, and targeted neurons were approached with the electrode. When near the neuron (small dent forming in neuron), positive pressure was released and a gigaohm (GΩ) seal (giga-seal) was formed between the tip of the electrode and the cell membrane. Gentle pulses of negative pressure were used to break through the membrane. Membrane holding potential was set to –65 mV and both fast and slow capacitance components were compensated for. Series resistance was routinely compensated for at 80%. The neuron was held at -65 mV until it was time to introduce stimulation protocols. Values reported are not liquid junction potential corrected. pMN identity was confirmed post hoc by assessing axon projection patterns facilitated by the addition of sulforhodamine B to the intracellular recording solution (0.1%; Millipore Sigma). Electrical activity was amplified and filtered at 10 kHz or 20 kHz with a Multiclamp 700B (Axon Instruments, Molecular Devices). The data was then digitized with a Digidata 1550 (Molecular Devices). A HumBug Noise Eliminator (Quest Scientific) was used to attenuate 50/60 Hz electrical noise.

### M-current relaxation protocol

To reveal the electrophysiological signature of the M-current, we implemented the standard M-current relaxation protocol (Sharples et al., 2023; Verneuil et al., 2020). This consisted of holding the neuron at -10 mV and introducing a series of hyperpolarizing voltage steps lasting 1 second each. As the membrane potential becomes hyperpolarized, the M-current deactivates. Current responses show the resulting loss of outward current caused by M-current deactivation. From this, we can estimate the amplitude of the M-current by taking the difference between the peak of the current response and the current at steady-state at the end of the step.

### Basal membrane properties

To measure membrane properties, 10 repetitions of a 50 ms current step of -5 pA were applied to neurons that were held at -65 mV. The average drop in voltage in response to the current step was used to calculate input resistance, whole-cell capacitance, and the membrane time constant.

### Firing properties

To measure firing properties, a series of 5 pA current steps from 0 pA to 290 pA were applied to neurons that were held at -65 mV. Rheobase was determined as the first current step to elicit an action potential. Using the action potential generated at rheobase, spike threshold was set as the voltage at which the voltage response reached a slope of 10 mV/ms (Sharples et al., 2023).

### Pharmacology

For pharmacological inhibition and activation of Kv7.2/7.3 channels, 10 μM XE-991 (X2254, Millipore Sigma) and 10 μM ICA-069673 (SML1616, Millipore Sigma) were first dissolved in DMSO (D8418, Millipore Sigma) before final dissolution in aCSF of the recording bath. XE-991 and ICA-069673 were introduced for a minimum of 10 minutes before proceeding with stimulation protocols for data collection.

### Data analysis and statistical analysis

All electrophysiological data was saved as .abf files. We used the open-source pyABF python package to import and read .abf files in Spyder (version 5.1.5). Analysis of recordings was semi-automated using Python (version 3.9.12) scripts tailored to each type of recording. Statistical analysis was performed using Prism by GraphPad (Version 10.3.1 (464)). When comparing the means of two non-normally distributed matched paired data sets, the non-parametric Wilcoxon matched-pairs signed rank tests were used. For unmatched pairs, Mann-Whitney tests were used. For comparisons between 3 data sets or more, one-way ANOVA with Tukey’s multiple comparisons test was used for normally distributed data and Kruskal-Wallis test with Dunn’s multiple comparisons test was used for non-parametric data. For all tests, significance stars are displayed on graphs.

## RESULTS

### M-current sensitive to Kv7.2/7.3 channel inhibitor XE-991 revealed in pMNs

We first sought out to determine whether primary motoneurons (pMNs) in larval zebrafish express the (I_M_). To this end, we targeted pMNs for whole-cell patch-clamp electrophysiology in spinalized larval zebrafish aged 4 and 5 dpf (**Figure 1A**). To reveal the electrophysiological signature of I_M_, we introduced the standard I_M_ relaxation protocol under voltage-clamp mode. This consists of holding the neuron at -10 mV, a relatively depolarized membrane potential at which I_M_ should be active, and introducing a series of hyperpolarizing voltage steps from -10 mV to -80 mV in increments of 5 mV (**Figure 1B-C**). Current responses to this protocol revealed the electrophysiological signature of the I_M_ relaxation in pMNs. **Figure 1D** displays its current-voltage relationship (n = 27). In a different subset of pMNs, we reveal that the current is sensitive to XE-991, a Kv7.2/7.3 channel inhibitor, at a concentration of 10 μM (n = 11). The amplitude of I_M_ is significantly reduced over voltages ranging from -45 to -30 mV when 10 μM XE-991 is present (−30 mV: *p =* 0.0061; -35 mV: *p =* 0.0085; -40 mV: *p =* 0.0282; -45 mV: *p =* 0.0159, **Figure 1D**). Inhibition of I_M_ by 10 μM XE-991 is also revealed when comparing the peak amplitude of the current (control: mean = 18.9 +/-8.7 pA; XE-991: mean = 11.2 +/-6.7 pA; Mann-Whitney test: *p =* 0.0085, **Figure 1F**). While the voltage at which I_M_ is activated (determined for individual neurons as the voltage at which I_M_ is not zero) does not differ significantly from control when XE-991 is present (control: -48.0 +/-5.9 mV; XE-991: -43.6 +/-6.7 mV; Mann-Whitney test: *p =* 0.0770, **Figure 1G**), the voltage at which I_M_ reaches half its maximal amplitude is significantly depolarized by XE-991 (control: -33.7 +/-6.0 mV and XE-991: -26.9 +/-7.7 mV; Mann-Whitney test: *p =* 0.0097, **Figure 1H**). Overall, these data demonstrate that XE-991-sensitive I_M_ is present in pMNs of 4-5 dpf zebrafish larvae.

**Figure 1.**
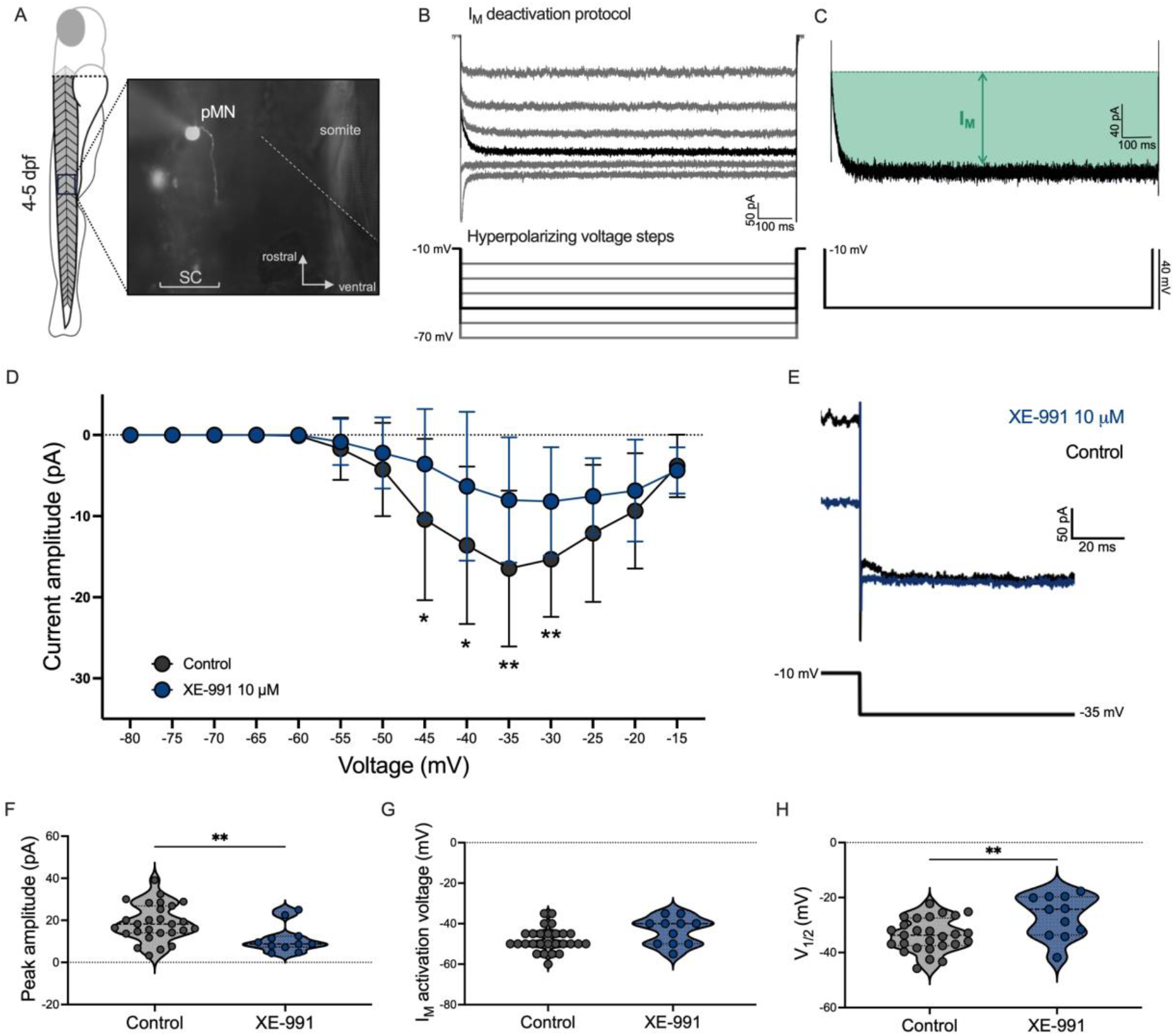
Whole-cell voltage-clamp reveals XE-991-sensitive I_M_ in primary motoneurons of larval zebrafish. **A**, Schematic depicting larvae aged 4-5 dpf with example of motoneuron filled with sulforhodamine B following patch-clamp recording in a 4 dpf larva. Dashed line demarcates the myotomal border between muscle somites. SC: spinal cord. pMN: primary motoneuron. **B**, *Top:* current traces in response to the standard I_M_ deactivation protocol during whole-cell recording. *Bottom:* command voltages consisting of stepping down from -10 mV to -70 mV in -5 mV voltage increments. **C**, Current trace response to a -40 mV step during the I_M_ deactivation protocol enlarged from **B** (black trace). The amplitude of I_M_ is estimated as the difference between the initial peak of the current and the steady-state current in response to the voltage step (green shaded area depicts I_M_ amplitude). **D**, Current-voltage (I-V) relationship of I_M_ measured in control pMNs (n = 26 cells from 11 fish; black) and during exposure to Kv7.2/7.3 channel inhibitor XE-991 (10 μM; n = 11 cells from 10 fish; blue). **E**, Example current traces from a pMN under control (black) and XE-991 treatment (blue) in response to a –25 mV hyperpolarizing voltage step during the I_M_ deactivation protocol. **F**, Peak I_M_ amplitude measured from pMNs under control (black) and XE-991 treatment (blue). **G**, Activation voltage of I_M_. **H**, Voltage at which I_M_ reaches half of its maximum amplitude. *Statistical analysis:* Mann-Whitney tests. * = *p* < 0.05, ** = *p* < 0.01; otherwise, not statistically significant.

### Pharmacological modulation of Kv7.2/7.3 channels alters pMN intrinsic properties

Having confirmed its presence in pMNs of larval zebrafish, we next investigated how I_M_ influences their intrinsic properties by pharmacologically modulating I_M_ conductance. Because I_M_ has been shown to be involved in setting resting membrane potential (Davis et al., 2020; Sharples et al., 2023; Verneuil et al., 2020; Wladyka and Kunze, 2006), we monitored the effects of pharmacologically activating then subsequently inhibiting Kv7.2/7.3 channels on resting membrane potential of pMNs (**Figure 2**). Whole-cell current-clamp recordings from pMNs of 4-5 dpf larvae reveal a significant hyperpolarization of mean resting membrane potential during Kv7.2/7.3 activation by 10 μM ICA-069673 (control (n = 10): -57.2 +/-2.5 mV; acute ICA-069673 (n =10): -67.7 +/-3.6 mV; prolonged ICA-069673 (n = 7): -70.6 +/-4.0 mV, **Figure 2A-B**). While inhibition of Kv7.2/7.3 by XE-991 (n = 7; -66.59 +/-3.8 mV) following activation by ICA-069673 did not depolarize pMN resting membrane potential to voltages near control levels, the slope describing the rate of change of voltage over time (control: -0.01 +/-0.01 mV/s; XE-991: 0.03 +/-0.01 mV/s; **Figure 2C**) as well as the change in membrane potential from start to end of each respective treatment (control: -1.2 +/-0.9 mV; XE 991: 6.4 +/-3.2 mV; **Figure 2D**) is significantly altered by XE-991 relative to control. These data demonstrate that I_M_ hyperpolarizes the membrane potential of pMNs.

**Figure 2.**
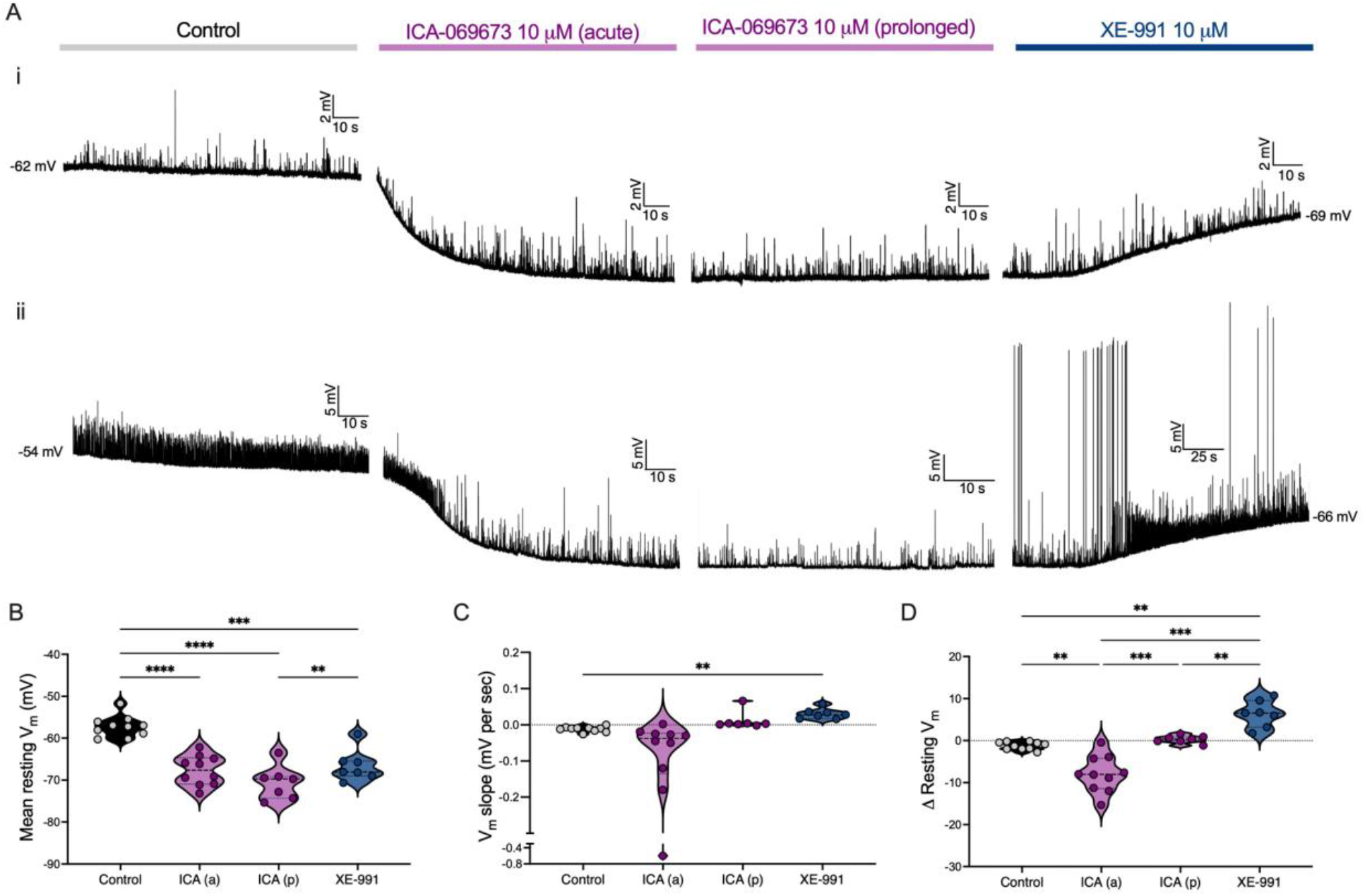
Pharmacological manipulation of Kv7.2/7.3 channels mediating I_M_ alters resting membrane potential of primary motoneurons. **A (i-ii)**, Whole-cell current-clamp traces from two pMNs with no holding current before (control; grey), during acute (under 3 minutes) and prolonged (10-15 minutes) exposure to Kv7.2/7.3 channel enhancer ICA-069673 (10 μM; magenta), and during subsequent exposure to XE-991 (10-15 minutes; 10 μM; blue). **B**, Mean resting membrane potential of pMNs during each treatment. **C**, Mean slope of membrane potential during current-clamp recordings of pMNs in each treatment. **D**, Overall change in resting membrane potential from start to finish of each treatment recording time. *Statistical analysis*, One-way ANOVA followed by Tukey’s multiple comparisons tests. ** = *p* < 0.01, *** = *p* < 0.001, **** = *p* < 0.0001; otherwise, not statistically significant.

We next sought out to evaluate how intrinsic firing and membrane properties of pMNs were affected by pharmacological modulation of I_M_. In the same group of pMNs, we held them at -65 mV before introducing 50 ms current steps in increments of 5 pA up to 195 pA, before (control) and after either Kv7.2/7.3 channel inhibition by 10 μM XE-991 or Kv7.2/7.3 channel activation by 10 μM ICA-069673 (**Figure 3**). Rheobase, spike threshold, and maximum number of spikes generated during the current steps were determined from these voltage responses (**Figure 3A-L**). While inhibition of I_M_ by XE-991 did not significantly affect rheobase of pMNs (*p =* 0.0557; n = 12; control: 91.3 +/-38.0 pA; XE-991: 85.4 +/-40.9 pA; **Figure 3C**), it significantly hyperpolarized spike threshold (*p =* 0.0034; n = 12; control: -29.9 +/-4.3 mV; XE-991: -33.3 +/-5.3 mV; **Figure 3D**) and increased the maximum number of spikes fired during the series of positive current steps (*p =* 0.002; n = 11; control: 2.7 +/-1.0 spikes; XE-991: 5.3 +/-2.2 spikes; **Figure 3E**). On the other hand, activation of I_M_ by ICA-069673 did not result in any significant changes to rheobase (*p =* 0.1021; n = 24; control: 121.5 +/-43.2 pA; ICA-069673: 111.5 +/-51.0 pA; **Figure 3F**), spike threshold (*p =* 0.4061; n = 24; -30.7 +/-6.2 mV; ICA-069673: -29. +/-6.78 mV; **Figure 3G**), or maximum number of spikes generated (*p =* 0.1759; n = 23; control: 2.4 +/-1.1 spikes; ICA: 3.0 +/-2.0 spikes; **Figure 3H**). Next, we introduced a 5 pA negative current step to pMNs held at -65 mV. From corresponding voltage responses, we calculated input resistance, whole-cell capacitance, and the membrane time constant (tau), before and after pharmacological modulation of I_M_. Our data reveal that inhibition of I_M_ by XE-991 does not alter input resistance (*p =* 0.5469; n = 8; control: 202.4 +/-71.2 MOhms; XE-991: 232.4 +/-101.6 MOhms; **Figure 3N**), whole-cell capacitance (*p =* 0.7422; n = 8; control: 22.1 +/-2.7 pF; XE-991: 22.5 +/-7.4 pF; **Figure 3O**), nor does it alter tau (*p =* 0.0781; n = 4.5 +/-1.8 ms; XE-991: 5.2 +/-2.3 ms; **Figure 3P**). In the presence of ICA-069673, input resistance is significantly increased (*p =* 0.0391; n = 8; control: 161.4 +/-40.3 MOhms; ICA-069673: 193.3 +/-58.4 MOhms; **Figure 3R**), but neither whole-cell capacitance (p > 0.9999; n = 8; control: 22.7 +/-5.4 pF; ICA-069673: 22.8 +/-3.4 pF; **Figure 3S**) nor tau (*p =* 0.0547; n = 8; control: 3.6 +/-0.9 ms; ICA-069673: 4.4 +/-1.5 ms; **Figure 3T**) are affected. Overall, these data demonstrate a role for I_M_ in regulating firing properties of pMNs in zebrafish larvae.

**Figure 3.**
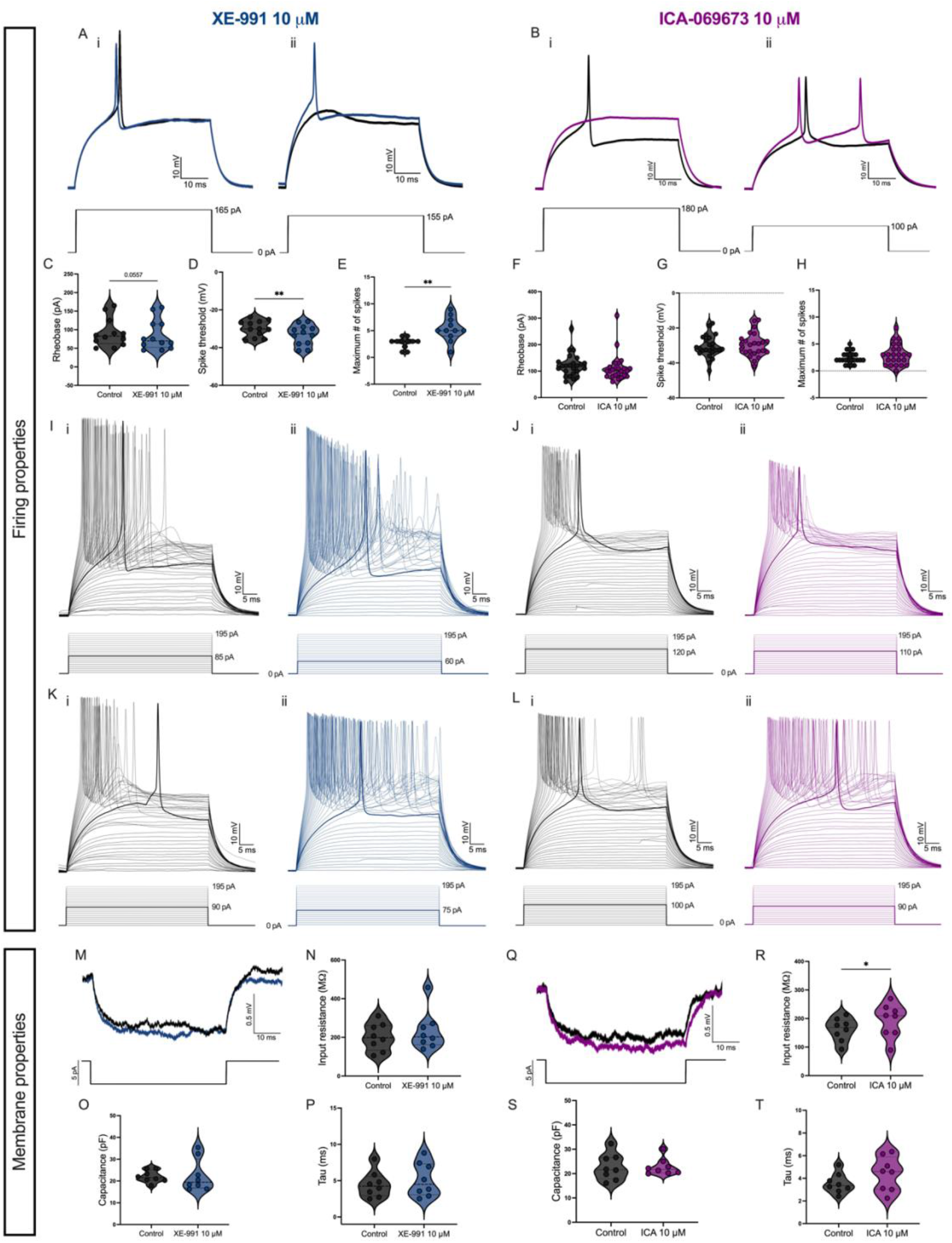
Pharmacological inhibition of Kv7.2/7.3 channels mediating I_M_ by XE-991 increases excitability of primary motoneurons in 4-5 dpf larval zebrafish. **A**, Examples of action potentials generated by positive current steps in the same pMN before (black) and after 10 μM XE-991 (blue) during whole-cell current-clamp recording in 4 dpf larva. Left and right panels (i-ii) show traces obtained from two pMNs. **B**, Examples of action potentials generated by positive current steps in the same pMN before (black) and after 10 μM ICA-069673 (magenta). Left and right panels (i-ii) show traces obtained from two pMNs. **C**, Rheobase before and after 10 μM XE-991. **D**, Spike threshold before and after 10 μM XE-991. **E**, Maximum number of spikes generated during a series of 50 ms current steps from 0 pA to 195 pA before (black) and after 10 μM XE-991 (blue). **F**, Rheobase before and after 10 μM ICA-069673. **G**, Spike threshold before and after 10 μM ICA-069673. **H**, Maximum number of spikes generated during a series of 50 ms current steps from 0 pA to 195 pA before (black) and after 10 μM ICA-069673 (magenta). **I, K**, Example voltage traces from two pMNs in response to a series of 50 ms current steps from 0 pA to 195 pA before (i, black) and after 10 μM XE-991 (ii, blue). **J, L**, Example voltage traces from two pMNs in response to a series of 50 ms current steps from 0 pA to 195 pA before (i, black) and after 10 μM ICA-069673 (ii, magenta). **M**, Example voltage trace from the same pMN in response to a –5 pA current step before (black) and after 10 μM XE-991 (blue). **N**, Input resistance before and after XE-991. **O**, Whole-cell capacitance before and after 10 μM XE-991. **P**, Membrane time constant (tau) before and after 10 μM XE-991. **Q**, Example voltage trace from the same pMN in response to a –5 pA current step before (black) and after 10 μM ICA-069673 (blue). **R**, Input resistance before and after 10 μM ICA-069673. **S**, Whole-cell capacitance before and after 10 μM ICA-069673. **T**, Membrane time constant (tau) before and after ICA-069673. *Statistical analysis*, Wilcoxon matched-pairs signed rank tests: * = p < 0.05, ** = p < 0.01; otherwise, not statistically significant.

### Developmental differences in the magnitude of I_M_ in pMNs

With evidence to support a role for I_M_ in regulating excitability of pMNs in larval zebrafish (**Figure 2**), we hypothesized that this role might change throughout development. As slow, low amplitude swimming movements start to dominate, the involvement of pMNs in locomotion diminishes. We proceeded to characterize properties of I_M_ in pMNs of larvae aged 2 dpf to 5 dpf (**Figure 4**). At 2 dpf, larvae still display coiling and can perform burst swimming in response to touch. By 3 dpf, they start to perform spontaneous burst swimming and by 4 dpf onward, larval zebrafish perform refined swimming known as beat-and-glide swimming. We had hypothesized that as zebrafish spinal locomotor circuits mature to generate increasingly refined movements relying less and less on pMNs, increasingly more dampening of intrinsic excitability by I_M_ may be necessary to ensure pMNs are active only during the fastest movements (e.g. escapes). We did indeed find that the amplitude of I_M_ from 2 dpf to 3 dpf triples (*p =* 0.0001; 2 dpf (n = 10): 9.7 +/-6.5 pA; 3 dpf (n = 11): 32.6 +/-14.6 pA; **Figure 4A-C**). Subsequently, I_M_ is reduced by nearly half by 4 dpf (*p =* 0.0476; 4 dpf (n = 16): 17.5 +/-8.5 pA; **Figure 4A-C**). From 4 dpf to 5 dpf, there is no change in peak I_M_ amplitude (p > 0.9999; 5 dpf (n = 11): 21.0 +/-9.0 pA; **Figure 4A-C**). When comparing voltage-dependent properties, our data show that the activation voltage of I_M_ is significantly depolarized at 2 dpf compared to 3 dpf (*p =* 0.0001; 2 dpf (n = 10): -36.0 +/-9.9 mV; 3 dpf (n = 11): -53.2 +/-4.6 mV; **Figure 4D**). Otherwise, no significant difference in the activation voltage of I_M_ is found between the age groups (4 dpf (n = 16): -47.2 +/-6.8 mV; 5 dpf (n = 11): -49.1 +/-4.4 mV; **Figure 4D**). Furthermore, we find that the voltage at which I_M_ is activated at half its maximal amplitude is significantly depolarized at 2 dpf (n = 9; -19.4 +/-7.3 mV) compared to 3 dpf (p < 0.0001; 3 dpf (n = 11): -37.4 +/-3.2 mV), 4 dpf (*p =* 0.0368; 4 dpf (n = 16): -32.1 +/-5.9 mV), and 5 dpf (*p =* 0.0019; 5 dpf (n = 11): -35.9 +/-5.5 mV; **Figure 4E**). These data demonstrate that both amplitude and voltage-dependent properties of I_M_ change during the 2 dpf to 5 dpf developmental time window.

**Figure 4.**
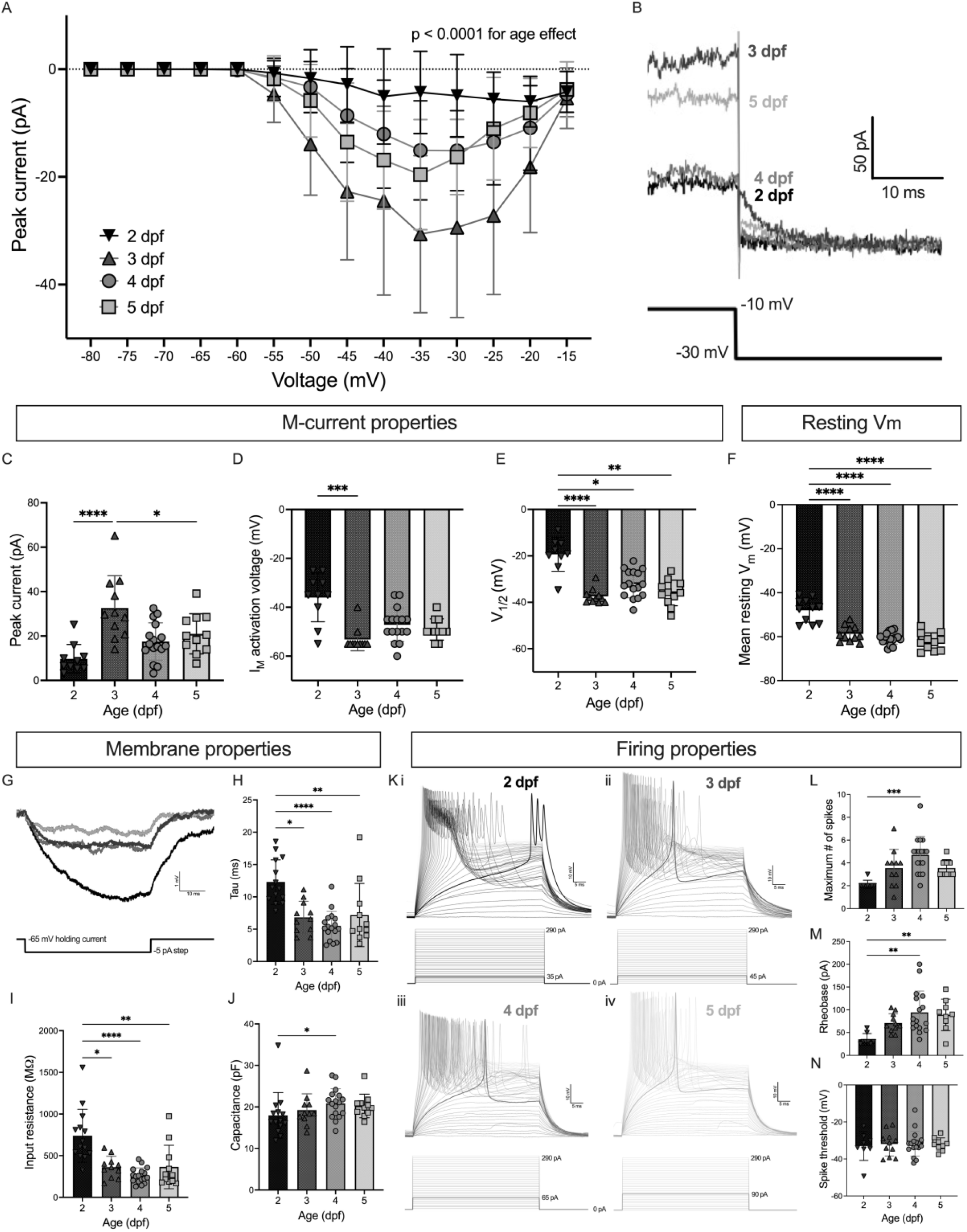
Differences in I_M_ in primary motoneurons at different ages of developing zebrafish. **A**, Current-voltage (I-V) relationship of I_M_ measured in pMNs from 2-5 dpf zebrafish. **B**, Example current traces showing I_M_ deactivation in 2-5 dpf pMNs in response to a –20 mV voltsage step during the standard I_M_ deactivation protocol. Comparison of **C**, peak I_M_, **D**, activation voltage of I_M_, E, voltage at which the I_M_ reaches half of its maximum amplitude, and **F**, mean resting membrane potential of 2-5 dpf pMNs. **G**, Example voltage traces in response to a –5 pA current step from pMNs of zebrafish aged 2 (black), 3 (dark grey), 4 (grey), and 5 dpf (light grey). Comparison of **H**, membrane time constant. **I**, input resistance. **J**, capacitance of 2-5 dpf pMNs. **K**, Example voltage traces in response to a series of positive 5 pA current steps from 0 pA to 290 pA in pMNs from larvae aged 2 (i), 3 (ii), 4 (iii), and 5 (iv) dpf. Comparison of: **L**, maximum number of spikes generated in response to a 290 pA current step; **M**, rheobase; **N**, spike threshold of 2-5 dpf pMNs. *Statistical analysis*, **B**: Effect of age assessed via mixed effects model. **C-F, H-J, & L-N:** Kruskal-Wallis test followed by Dunn’s test for multiple comparisons. * = *p* < 0.05, ** = *p* < 0.01, *** = *p* < 0.001, **** = p < 0.0001; otherwise, not statistically significant.

Next, we sought to determine whether differences in magnitude and voltage-dependency of I_M_ in pMNs across ages coincided with any differences in intrinsic properties of the pMNs throughout development. We first compared the mean resting membrane potential of pMNs across the four age groups. The membrane potential at rest in 2 dpf pMNs was significantly depolarized compared to the three older age groups (p < 0.0001; 2 dpf (n = 11): -48.0+/-5.4 mV; 3 dpf (n = 11): -58.7 +/-3.6 mV; 4 dpf (n = 16): -60.7 +/-2.7 mV; 5 dpf (n = 11): -62.0 +/-3.9 mV; **Figure 4F**). This relatively depolarized resting membrane potential at 2 dpf coincides with a relative reduction in the magnitude of I_M_ at 2 dpf (**Figure 2A-C**).

pMNs at 2 dpf displayed a significantly higher input resistance compared to those at 3 dpf (*p =* 0.0341; 2 dpf (n = 14): 738.4 +/-317.8 MOhms; 3 dpf (n = 11): 362.0 +/-131.1 MOhms), 4 dpf (p < 0.0001; 4 dpf (n = 17): 261.9 +/-91.9 MOhms), and 5 dpf (*p =* 0.0030; 5 dpf (n = 11): 364.4 +/-261.1 MOhms; **Figure 4I**). Similarly, 2 dpf pMNs display higher values of the membrane time constant (tau), compared to all other ages (2 dpf versus 3 dpf: *p =* 0.0174; 2 dpf versus 4 dpf: p < 0.0001; 2 dpf versus 5 dpf: *p =* 0.0058; 3 dpf versus 4 dpf: p > 0.9999; 3 dpf versus 5 dpf : p > 0.9999; 4 dpf versus 5 dpf > 0.9999; 2 dpf (n = 14): 12.3 +/-3.4 ms; 3 dpf (n = 11): 6.8 +/-2.5 ms; 4 dpf (n = 17): 5.5 +/-2.3 ms; 5 dpf (n = 11): 7.2 +/-4.9 ms; **Figure 4H**). Whole-cell capacitance on the other hand shows a significant increase only from 2 dpf to 4 dpf (2 dpf versus 3 dpf: p > 0.9999; 2 dpf versus 4 dpf: *p =* 0.0262; 2 dpf versus 5 dpf: *p =* 0.1841; 3 versus 4 dpf: *p =* 0.8446; 3 dpf versus 5 dpf: p > 0.9999; 4 dpf versus 5 dpf: p > 0.9999; 2 dpf (n = 14): 18.0 +/-5.5 pF; 3 dpf (n = 11): 19.2 +/-3.9 pF; 4 dpf (n = 17): 20.9 +/-3.6 pF; 5 dpf (n = 11): 20.2 +/-2.8 pF; **Figure 4J**).

Finally, we investigated changes in firing properties of pMNs from 2 to 5 dpf (**Figure 4K-N**). There was a significant increase in the number of spikes generated in response to a maximal applied current (290 pA), 50 ms current step from 2 dpf to 4 dpf (*p =* 0.0001) but no other significant differences were observed across the ages (2 dpf vs 3 dpf: *p =* 0.1214, 2 dpf vs 5 dpf: *p =* 0.1044, 3 dpf vs 4 dpf: *p =* 0.3378, 3 dpf vs 5 dpf: *p* > 0.9999, 4 dpf vs 5 dpf: *p =* 0.6498; 2 dpf (n = 8): 2.125 +/-0.3536 spikes; 3 dpf (n = 11): 3.5 +/-1.6 spikes; 4 dpf (n = 17): 4.7 +/-1.6 spikes; 5 dpf (n = 9): 3.6 +/-0.7 spikes; **Figure 4L**). Rheobase was significantly reduced in pMNs at 2 dpf compared to 4 dpf and 5 dpf, but not 3 dpf (2 dpf vs 3 dpf: *p =* 0.0614, 2 dpf vs 4 dpf: *p =* 0.0010, 2 dpf vs 5 dpf: *p =* 0.0034, 3 dpf vs 4 dpf: *p* > 0.9999, 3 dpf vs 5 dpf: *p* > 0.9999, 4 dpf vs 5 dpf: *p* > 0.9999; 2 dpf: 35.0 +/-12.8 pA; 3 dpf: 70.9 +/-20.6 pA; 4 dpf: 94.1 +/-47.1; 5 dpf: 88.9 +/-34.5 pA; **Figure 4M**), which is in line with the observed increase in input resistance at 2 dpf. No significant differences in spike threshold were found across ages (for all comparisons: *p* > 0.9999; 2 dpf: -33.0 +/-7.8 mV; 3 dpf: -31.9 +/-6.5 mV; 4 dpf: -31.8 +/-6.7; 5 dpf: -32.1 +/-3.5 mV; **Figure 4N**).

Overall, these data reveal a transient but important increase in the magnitude of I_M_ in 3 dpf pMNs. They also reveal important differences in intrinsic properties notably between 2 dpf and older pMNs. While the amplitude of I_M_ is significantly larger at 3 dpf compared to 4 dpf, there are no accompanying changes to pMN firing and intrinsic properties.

## DISCUSSION

Since its initial discovery (Brown and Adams, 1980), the M-current has been shown to limit neuronal excitability and burst firing observed in neurons in many systems (Adams, Paul R., Halliwell, James V., 1982; Adams et al., 1982; Bordas et al., 2015; Davis et al., 2020; Drion et al., 2010; Santini and Porter, 2010). The presence of the M-current in spinal locomotor circuit neurons was recently demonstrated, where it works in tandem with the persistent sodium current (I_NaP_) to control burst firing in spinal neurons, ultimately exerting an important control over locomotor speed (Verneuil et al., 2020). Recently, Sharples et al. showed that varying magnitudes of the M-current are expressed in subtypes of mouse motoneurons as a means to control their recruitment (Sharples et al., 2023). Until now, whether the M-current was expressed in larval zebrafish motoneurons and involved in controlling their excitability remained uninvestigated.

We used patch-clamp electrophysiology in combination with pharmacology to reveal, for the first time, the presence of a XE-991 sensitive M-current in primary motoneurons. Inhibition by XE-991 or activation by ICA-069673 modified resting membrane potentials, which is consistent with the subthreshold nature of the M-current. We observed that inhibition of the M-current by XE-991 altered firing properties of pMNs, but activation by ICA-069673 did not. Specifically, enhancing the M-current with ICA-069673 altered none of the measures of excitability we measured (rheobase, spike threshold, and number of spikes generated) despite observing that ICA-069673 significantly hyperpolarized resting membrane potential, consistent with its effects in other studies (Davis et al., 2020; Sharples et al., 2023; Verneuil et al., 2020; Wladyka and Kunze, 2006). It is possible therefore that the baseline level of the M-current in pMNs is near maximal levels, or activating the M-current is affecting other components of the locomotor circuits, altering synaptic activity at the level of the pMNs, and ultimately affecting membrane dynamics of pMNs such that effects to firing behaviours are masked.

### Transient increase of the M-current in pMNs at 3 dpf

Considering the presence of the M-current in larval pMNs, we then set out to understand how membrane dynamics of pMNs might change throughout development as a means to foster proper recruitment of motoneurons as locomotor movements that do not rely on pMNs become more prominent. We hypothesized that if the M-current were present in zebrafish pMNs, we might observe the magnitude of the M-current increasing from 2 dpf to 5 dpf, as the need to limit recruitment of pMNs increases as slow swimming movements emerge. What we find instead is a transient increase in the amplitude of the M-current at 3 dpf, tripling from its peak amplitude at 2 dpf and subsequently getting reduced to almost half its 3 dpf amplitude by 4 dpf.

One possible explanation for the increase in M-current magnitude from 2 dpf to 3 dpf is an increase in the expression of ion channels that mediate the M-current. An increase in the expression of Kv7.2/7.3 channels at 3 dpf could underlie an increase in the magnitude of observed M-current in pMNs at this age. Single cell transcriptional data from developing zebrafish from the open-source database, *Daniocell*, suggest that the *KCNQ3* gene encoding the Kv7.3 subunit may indeed be more highly expressed in motoneurons at 3 dpf compared to 2 dpf, though the differences are subtle (Sur Abhinav et al., 2023). Alternatively, a change in the conductance of the M-current could explain the differences in magnitude. Previous work has identified phosphatidylinositol 4,5-bisphosphate (PIP_2_) as an important regulator of Kv7.2/7.3 channel activity whereby opening of Kv7.2/7.3 channels requires membrane PIP_2_ (Li et al., 2005; Zhang et al., 2003). It is therefore possible that increases in levels of PIP_2_ from 2 dpf to 3 dpf, brought on either by upregulation of PIP_2_ synthesis or downregulation of PIP_2_ breakdown enzymes, for example, underlie an increase in Kv7.2/7.3 activity. Experiments directly linking Kv7.2/7.3 expressional data to pMNs would be useful to delineate the mechanism underlying the increase of the M-current at 3 dpf from 2 dpf. This would be further complimented by investigation into the expression of PIP_2_ in pMNs across development.

Whichever is the underlying mechanism, the observed increase of the M-current at 3 dpf was transient, diminishing by nearly half its maximal amplitude by 4 dpf. While changes to Kv7.2/7.3 expression and conductance levels similar to those described above may also underlie the apparent reduction in M-current by 4 dpf, another possibility is the involvement of other ionic conductances to pMN membrane dynamics. While the standard M-current relaxation protocol used in this study reveals the electrophysiological signature of M-current deactivation, the resulting current responses do not exclude voltage-gated inward currents. Persistent inward sodium currents (I_NaP_) mediated by ion channels from the Na_V_ family activate with depolarization of membrane potential (Crill, 1996; Raman and Bean, 2001). We cannot exclude the possibility that deactivation of a Na_V_-mediated inward current during hyperpolarizing voltage steps of the M-current relaxation protocol might mask the true magnitude of the M-current. Electrophysiology and/or Na_v_ ion channel expression experiments investigating the presence of I_NaP_ in pMNs would be critical in delineating the cause of the reduction in M-current magnitude from 3 dpf to 4 dpf.

If there is no masking of the M-current by I_NaP_ in our recordings at 4 and 5 dpf, and there is a true decrease in the M-current after 3 dpf, then our findings demonstrating changes in basal membrane and firing properties of pMNs only from 2 dpf to 3 dpf cannot be explained solely by the M-current. The lack of difference in resting membrane potential of 3 dpf pMNs compared to 4 dpf and 5 dpf pMNs despite a decrease in the M-current could instead be due to a parallel increase in some other outward current and/or a decrease in some inward ion current to stabilize resting membrane potential.

Overall, these data suggest that the influence of the M-current on basal excitability of pMNs peaks at 3 dpf, whether because the amplitude of this current is greatest at that stage or other ionic currents start to take on a more prominent role in regulating pMN membrane dynamics together with the M-current by 4 dpf.

### The M-current and behaviour

The consequences of the possible transient peak of the M-current in pMNs around 3 dpf for motor control in developing zebrafish remains to be investigated. pMNs are known to be necessary for the execution of the fastest swims, such as the escape response, in larval zebrafish (Fetcho and O’Malley, 1995; Liu and Westerfield, 1988). Our data show that pharmacological inhibition of the M-current increases excitability of pMNs and promotes the generation of more spikes during a given current injection. It is possible that the M-current controls the recruitment of pMNs, limiting their involvement to large amplitude and fast movements where constraining the number of action potentials generated fosters quicker coordination of muscle contraction (Verneuil et al., 2020). Furthermore, the control of locomotion by spinal circuits in zebrafish is speed-dependent, with speed-specific spinal circuits controlling slow, intermediate, and fast swimming (Ampatzis et al., 2014; McLean et al., 2007). The M-current may act to restrict the activation of pMNs during slower locomotion and ensure that pMNs are only active upon sufficient synaptic excitation when necessary, as locomotor speeds increase.

### The M-current in spinal locomotor circuits

We have focused our initial investigation of the M-current in larval zebrafish spinal neurons to pMNs. Given the recent evidence demonstrating that subtypes of mouse motoneurons express varying magnitudes of the M-current to control their recruitment (Sharples et al., 2023), secondary motoneurons (sMNs) involved in lower frequency swimming (McLean et al., 2007) may also express the M-current. Should this be the case, whether their respective M-current magnitudes are distinct such that they reflect their differences in firing behaviours (Menelaou and McLean, 2012) will be useful to understand the relative contribution of the M-current in setting up membrane dynamics suited for different firing behaviours.

Verneuil et al. demonstrated for the first time a role for the M-current in locomotion, showing that Kv7.2/7.3 ion channel expression is ubiquitous across spinal neurons for locomotion and helps control neuronal bursting in rodents (Verneuil et al., 2020). Investigation into the identities of larval zebrafish spinal neurons, other than motoneurons, that might express the M-current will elucidate similarities in the role the M-current plays in spinal circuits for locomotion across vertebrate species.

### Kv7.2/7.3 channels as targets of neuromodulation

Ion channels have been shown to be important targets of neuromodulation because their activity guides neuronal membrane and firing properties. In the case of spinal neurons, neuromodulation of ion channels by neurotransmitters such as serotonin or dopamine is linked to changes in motor activity (Guertin and Hounsgaard, 1998; Hounsgaard and Kiehn, 1985; Perrier et al., 2003; Picton and Sillar, 2016). The M-current gets its name from the finding that it is inhibited by acetylcholine via muscarinic receptors in frog sympathetic neurons (Brown and Adams, 1980). Whether the M-current in pMNs is similarly modulated by acetylcholine in larval zebrafish remains unknown. As spinal locomotor circuits are developing, so is the establishment of supraspinal descending neuromodulatory systems and their influence on these circuits (Brustein et al., 2003; Lambert et al., 2012; Thirumalai and Cline, 2008). While the relative contribution of the M-current to basal excitability of pMNs seems to taper off after 3 dpf, the continued presence of this current presents a lever for descending systems to shape the excitability of pMNs. Revealing the M-current in pMNs and changes in its amplitude and voltage-dependency in early zebrafish development is an important step to understanding the contributions of this current to motor control. Identifying which neuromodulators and their receptors might modulate the activity of the M-current will further shed light onto how the activity of spinal locomotor circuits is fine-tuned supraspinally, either as a result of development or depending on environmental contexts.

To conclude, our data reveal a novel ionic current expressed in primary motoneurons of larval zebrafish. We demonstrate that the control of motoneuron excitability observed in mice (Sharples et al., 2023) is conserved in larval zebrafish. What underlies the developmental changes in the magnitude of the M-current in pMNs as well as how this impacts the emergence and modulation of locomotor behaviour through the M-current will further reveal how the nervous system controls movements across vertebrates.

## Acknowledgements

This research was supported by the Natural Sciences and Engineering Research Council of Canada (NSERC) PGS-D 569969-2022, NSERC CGS-M 553401-2020, and NSERC Discovery Grant RGPIN-2015-06403.

## Competing interests

The authors declare no competing interests.

## Notes

### Competing Interest Statement

The authors have declared no competing interest.

